# Mice expressing P301S mutant human tau have deficits in interval timing

**DOI:** 10.1101/2022.04.04.487032

**Authors:** Travis Larson, Vaibhav Khadelwal, Matthew A. Weber, Mariah R. Leidinger, David K. Meyerholz, Nandakumar S. Narayanan, Qiang Zhang

## Abstract

Interval timing is a key executive process that involves estimating the duration of an interval over several seconds or minutes. Patients with Alzheimer’s disease (AD) have deficits in interval timing. Since temporal control of action is highly conserved across mammalian species, studying interval timing tasks in animal AD models may be relevant to human disease. Amyloid plaques and tau neurofibrillary tangles are hallmark features of AD. While rodent models of amyloid pathology are known to have interval timing impairments, to our knowledge, interval timing has not been studied in models of tauopathy. Here, we evaluate interval timing performance of P301S transgenic mice, a widely studied model of tauopathy that overexpresses human tau with the P301S mutation. We employed the switch interval timing task, and found that P301S mice consistently underestimated temporal intervals compared to wild-type controls, responding early in anticipation of the target interval. Our study indicating timing deficits in a mouse tauopathy model could have relevance to human tauopathies such as AD.

**HIGHLIGHTS:**

- We examined interval timing behavior in mice expressing P301S mutant tau
- P301S mice responded earler than littermate controls
- These data provide insight into animal models of tauopathy

## INTRODUCTION

Patients with Alzheimer’s disease (AD) have deficits in multiple cognitive domains, including long-term memory and executive functions (Marshall et al., 2011) such as working memory, attention, planning, and timing (Brown, 2006; Gilbert & Burgess, 2008). While the neurobiology of amnestic dysfunction has been previously studied (Jahn, 2013), impairment of executive function in AD is less understood. Although there are mouse models of key pathophysiological processes in AD (Kimura & Ohno, 2009; Yoshiyama et al., 2007), studying executive functions in mice can be challenging, because it is difficult to identify experimental paradigms with translational validity between rodents and humans. Identifying executive function deficits in AD mouse models is critical to developing novel interventions for these debilitating aspects of AD.

A paradigm that has been used to study executive function in rodent models is interval timing, which requires subjects to estimate an interval of several seconds with a motor response (Buhusi & Meck, 2005). Interval timing requires working memory to follow temporal rules and attention to the passage of time (Merchant & de Lafuente, 2014). Interval timing is an ideal executive function to study in translational models, as deficits in mice can have relevance for human diseases (Parker et al., 2013; Ward et al., 2011). Interval timing is impaired in AD patients, with greater variability and distortions in temporal processing relative to controls (Carmen Carrasco, 2000; Caselli et al., 2009; El Haj & Kapogiannis, 2016). These deficits may be predictive of early disease in AD (Bangert & Balota, 2012). Familial AD can be caused by mutations in the amyloid precursor protein and presenilin, both of which lead to aggregation of beta-amyloid plaques (Goedert & Spillantini, 2006). Rodent models of beta-amyloid aggregation, including APP/swe and 5xFAD, have deficits in interval timing (Armstrong et al., 2020; Gür, Fertan, Alkins, et al., 2019, respectively). However, AD can also involve intracellular aggregation of tau, a microtubule-associated protein (Lee et al., 2001; Schwarz et al., 2016). Abnormal tau is a feature of several brain disorders, including frontotemporal dementia and corticobasal degeneration (Yoshiyama et al., 2001). Rodent models of tauopathy have widespread synaptic and cognitive deficits (Yoshiyama et al., 2007), leading to our hypothesis that mice expressing mutant tau will have interval timing deficits.

We tested this hypothesis using a P301S transgenic mouse model (Yoshiyama et al., 2007). These transgenic mice possess the human P301S mutant form of tau, leading to a five-fold higher expression of tau-4R, one of the two major isoforms of tau (Rademakers et al., 2004). We trained these animals to perform a “switch” interval timing task, in which they must switch from one nosepoke to the other after a temporal interval to receive a reward (Balci et al., 2008; Bruce et al., 2021; Tosun et al., 2016). We found that P301S mice switch at earlier times compared to non-transgenic controls, producing a leftward shift in time-response functions. These data extend our understanding of executive dysfunction in animal models of tauopathy, which could be useful for developing new biomarkers or therapies for human taupathies, such as AD and frontotemporal dementia.

## MATERIAL AND METHODS

### Mice

Experimental procedures were approved by the Institutional Animal Care and Use Committee (IACUC) at the University of Iowa and performed in accordance with the guidelines set forth in Protocol #0062039. Thirteen female P301S transgenic mice, (Jackson Labs, Bar Harbor, ME; Strain #008169, founder line 19) and 9 wild-type female B6C3F1/J littermates were communally housed on a 12 hour light/dark cycle. Both the transgenic and non-transgenic mice began training for the switch interval timing task at approximately 6 months of age. Mice were weighed daily and kept on a restricted diet for the duration of the experiment. Water was available ad libitum.

### Switch interval timing task

The switch interval timing task (Fig. 1A) assesses an animal’s ability to control actions based on an internal representation of time (Balci et al., 2008; Bruce et al., 2021; Tosun et al., 2016). Mice were trained to perform the task in operant chambers (MedAssociates, St. Albans, VT) placed in sound attenuating cabinets. The chambers contained two response ports at the front, separated by a reward hopper, and one response port at the back, opposite the reward hopper. Cues were generated by a light located above each front port and a speaker that produced an 8-kHz tone at 72 dB during the trials. Infrared beams at each port were used to detect the mouse’s responses through nosepokes. A trial was initiated by a nosepoke at the back, following which the cue lights and tone turned on for either a 6-second short trial or an 18-second long trial. Short and long trials had identical cues. Short trials were reinforced with 20-mg sucrose pellets (BioServ, Flemington, NJ) for the first response after 6 seconds at the designated “short port” (front-left or front-right, counterbalanced across mice). Long trials were reinforced only when the mouse responded after 18 seconds by “switching” from the short port to the long port. On these switch trials, mice responded at the short port until after 6 seconds, when they switched to responding at the long port until reward delivery. Once trained for the switch interval timing task, four sessions of test data per mouse were collected and analyzed. Experimental sessions lasted 90 minutes and consisted of equal numbers of short and long trials. Only data from long trials were analyzed. We did not analyze data from two mice (one control and one P301S) that did not complete more than five switch trials.

**Figure 1:**
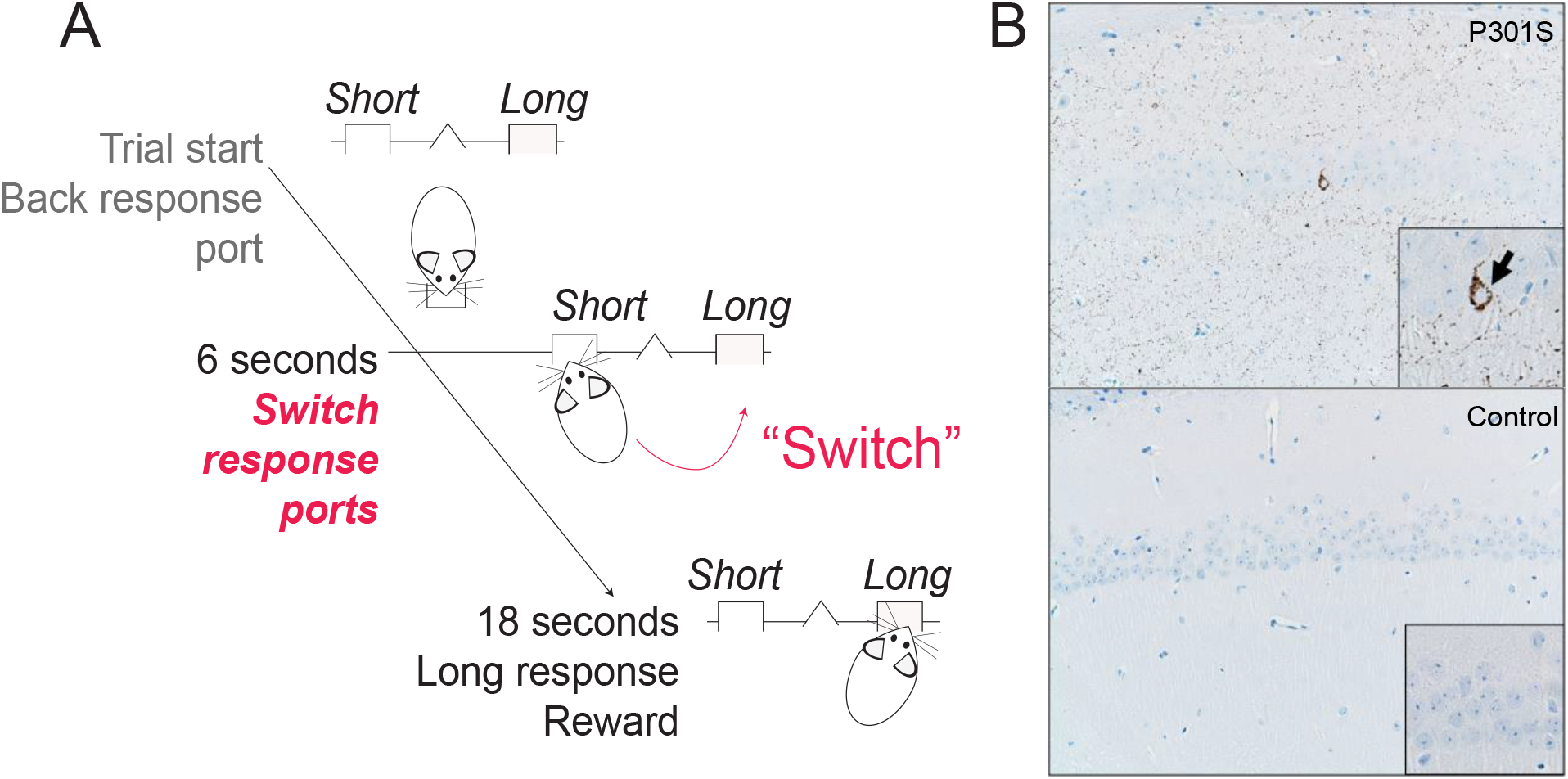
Switch interval timing task and hippocampal tau pathology in P301S transgenic mice. **A)** In the switch interval timing task, mice initiate trials at a back response port. On half the trials, mice were rewarded at the designated short response port after 6 seconds; for the remaining trials, mice were rewarded at the long response port after 18 seconds. The temporal decision to switch (in red) from the short to the long response port is an explicit time-based decision, as in other interval timing tasks. We focused our analysis on these long “switch” trials. **B)** P301S mice have pathological tau accumulation. Phospho-tau antibody (AT8; in brown) staining showed robust phospho-tau deposits in the hippocampus (CA1) of P301S mice. No significant tau deposits were visible in the non-transgenic controls.

### Immunohistochemistry

At approximately 8 months of age, mice were anesthetized with isofluorane and transcardially perfused with ice-cold phosphate-buffered saline. Brains were collected and postfixed in 4% paraformaldehyde overnight followed by 30% sucrose for approximately 48 hours. The Comparative Pathology Laboratory at the University of Iowa performed all sectioning and immunohistochemistry to visualize tau pathology in the mouse brain (Hefti et al., 2019). Sections (4–5 µm) were collected and embedded in paraffin. Antigen retrieval was done using citrate buffer (pH 6.0) in the New Decloaker, 110 °C for 15 minutes, followed by incubation in phospho-tau monoclonal antibody AT8 (Thermo Fisher, MN1020, Waltham, Massachusetts, U.S.) antibody (1:1000 in Dako diluent buffer) for 15 minutes at room temperature. Sections were mounted, counterstained with hematoxylin for 3 minutes, and coverslipped. Slides were visualized using an Olympus BX53 microscope, DP73 digital camera, and CellSens Dimension Software (Olympus, Tokyo, Japan).

### Data Analysis

Data analysis was performed as described previously (Bruce et al., 2021). Our analysis was limited to long trials only from which we extracted the switch time of each trial, or the time after trial initiation. On swtich trials (Fig. 1A), the mouse made its last response at the short port, thereby capturing the time when the decision to switch was made. We quantified the cumulative probability of switch times, as well as the mean switch time and the coefficient of variation. Differences between groups were compared using the nonparametric Wilcoxon test. All statistical procedures were reviewed by the Biostatistics and Epidemiology Research and Design Core at the Institute for Clinical and Translational Sciences at the University of Iowa.

## RESULTS

### Tau pathology in the frontal cortex and hippocampus of P301S transgenic mice

We visualized the distribution of tau in the P301S mouse brain using immunohistochemical staining with the phospho-tau monoclonal AT8 antibody (Hefti et al., 2019). P301S mice showed strong tau positive immunostaining, particularly in the hippocampus (Fig. 1B). Tau pathology in the frontal cortex and striatum was also observed, but was less prominent. No significant tau deposits were apparent in comparable regions of non-transgenic control mice.

### P301S mice have earlier switch times

We trained 13 female P301S mice and 9 female littermate controls to perform the switch interval timing task. The distribution of switch times (Fig. 2A) indicates that both groups of mice tended to switch just after 6 seconds, the total duration of a short trial, implying successful acquisition of the task parameters. Additionally, the probability function for the transgenic mice was shifted to the left relative to the non-transgenic control mice, indicating that the P301S transgenic mice reliably switched earlier (median: 8.9 seconds (intraquartile range: 4.7–9.4 seconds)) than the non-transgenic control group (11.0 (10.5–11.2) seconds); Wilcoxon *p* < 0.001; Cohen’s *d* = 1.5; Fig. 2B). There was no reliable difference in the coefficient of variation between P301S and non-transgenic mice (Fig. 2C; P301S: 0.41 (0.36–0.44); non-transgenic: 0.35 (0.32-0.38); Wilcoxon *p* = 0.14; Cohen’s *d* = 0.75). There was also no reliable difference between the number of responses (P301S: 102 (75–126); non-transgenic: 95.25 (84–113); Wilcoxon *p* = 0.90; Cohen’s *d* = 0.14) and the total number of rewards (P301S: 40 (29–42); non-transgenic: 31 (27–34); Wilcoxon *p* = 0.08; Cohen’s *d* = 0.8).

**Figure 2:**
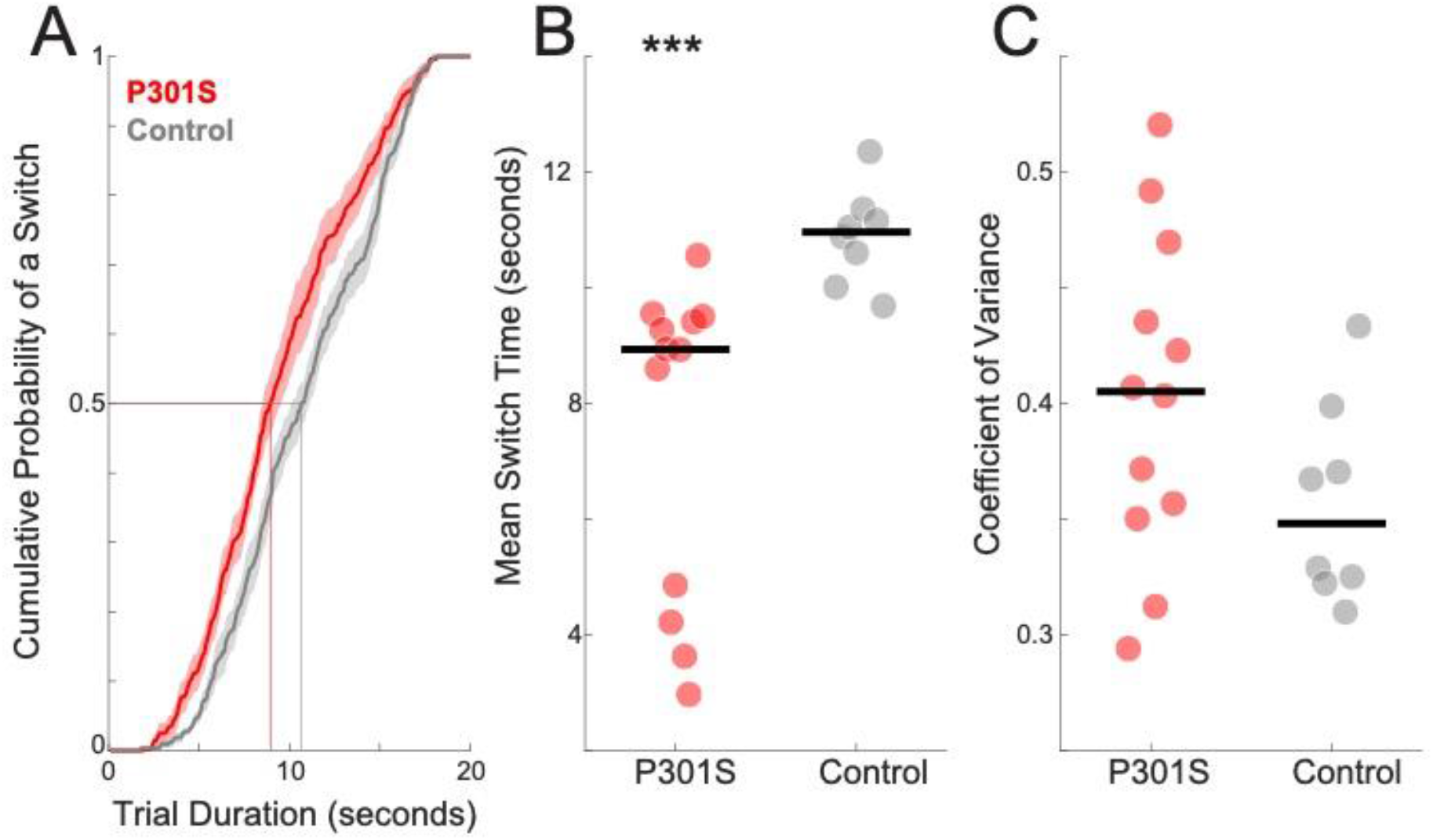
P301S transgenic mice reliably switched earlier than their non-transgenic counterparts. **A**) Cumulative probability distribution of switch times. **B)** Mean switch time and **C)** coefficient of variance across four 90-minute test sessions. Data from 12 P301S female mice (in red) and 8 non-transgenic control female mice (in gray). ***: *p* < 0.001.

## DISCUSSION

We examined interval timing performance in transgenic mice expressing human tau with the P301S mutation. We found consistent evidence of deficits in switch timing performance, with P301S mice switching earlier in the trial, without a change in variability or response rate. Our data indicates that animal tauopathy models can exhibit interval timing deficits consistent with anticipatory responding. Since multiple cognitive processes, such as motivation, working memory, and attention underlie temporal anticipatory behavior (Balsam et al., 2009), this insight could be useful both for a better understanding of tauopathies and for developing and testing new interventions aimed at executive dysfunction in human tauopathies.

Our work is supported by prior work on animal models of AD. Gür et al. (2019) found that 3xFAD mice had comparable interval timing performance to wild-type controls (Gür, Fertan, Kosel, et al., 2019). In a follow-up study in 5xFAD mice, the same group found that female 5xFAD mice responded significantly earlier with shorter peak times, consistent with increased anticipatory responding (Gür, Fertan, Alkins, et al., 2019), which might be associated with hippocampal-based memory processes. Similar results were found in the APPswe/PS1dE9 model of AD (Armstrong et al., 2020), with increased variability and earlier timing peaks. Our results from the switch interval timing paradigm also report an anticipatory “leftward” shift, with earlier switch times but no change in timing variability.

Both APPswe/PS1dE9 and 5xFAD mice have marked neurodegeneration of the hippocampus, as well as other structures, due to beta-amyloid pathology (Garcia-Alloza et al., 2006; Kimura & Ohno, 2009, respectively). Studies in rodents with hippocampal lesions using the peak interval procedure, in which subjects estimate a fixed time interval, have found similar leftward shifts in time estimation (Meck et al., 1984, 2013). The P301S tau model, which involves mutations in human tau, overlaps with frontotemporal dementia (Lee et al., 2001; Rademakers et al., 2004; Yoshiyama et al., 2007) and exhibits abormal tau in the frontal cortex and striatum, as well as the hippocampus (Yoshiyama et al., 2007). Because frontostriatal circuits play a key role in interval timing, they may have interacted with hippocampal deficits to produce the marked leftward shift we observed in this study (Emmons et al., 2020; Emmons et al., 2016; Meck et al., 1984).

Human AD patients have a range of hippocampal-dependent impairments (West et al., 1994). AD patients have decreased ability to discriminate between short intervals and have greater variance for temporal judgements in the sub-second range (Caselli et al., 2009). This variance carried through to longer intervals of 5–25 seconds (Carmen Carrasco, 2000). Such impairments could be exacerbated during dual-task conditions (Papagno et al., 2004) and might exist in patients with early AD (Bangert & Balota, 2012). Human AD is incredibly complex, involving multiple proteins, cellular processes, and brain regions; however, animals that model aspects of AD may be useful in developing new understanding of disease processes, biomarkers, and therapies. This study extends prior work on animal models of beta-amyloid to animal models of tau. Furthermore, the P301S transgenic mouse line models aspects of human frontotemporal dementia, which also can present with timing impairments (Wiener & Coslett, 2008).

Our study has several limitations. First, P301S mice overexpress tau-4R, one of the two major isoforms of tau, limiting our conclusions to only a specific tau isoform. Second, the distribution of accumulated tau is widespread, making it difficult for us to definitively trace the anatomical correlates of the observed differences. Future studies leveraging regional overexpression of tau might be able to localize tau overexpression to a specific circuit. Third, we restricted our study to female mice that were reported in prior work and were available from Jackson Labs (Armstrong et al., 2020). Finally, our use of interval timing may not capture the entire domain of cognitive deficits in human tauopathies.

Our study utilizes a mouse model of tauopathy and demonstrates that tau pathology alone might induce deficits in interval timing behavior. We observed high tau deposition in the hippocampus, reinforcing the importance of temporal memory in interval timing. We propose that future work should aim at generalizing these results to other models of tauopathy with targeted tau deposition, to better elucidate mechanisms and inform future clinical interventions.

